# Modeling methylation dynamics with simultaneous changes in CpG islands

**DOI:** 10.1101/638023

**Authors:** Konrad Grosser, Dirk Metzler

## Abstract

**Motivation:** Probabilistic models for methylation dynamics of CpG sites are usually based on sequence evolution models that assume indepedence between sites. In vertebrate genomes, CpG sites can be clustered in CpG islands, and the amount of methylation in a CpG island can change due to gene regulation processes. We propose a probabilistic model of methylation dynamics that accounts for simultaneous methylation changes in multiple CpG sites belonging to the same CpG island. We further propose a Markov-chain Monte-Carlo method to fit this model to methylation data from cell type phylogenies and apply this method to available data from murine haematopoietic cells.

**Results:** Branch lengths in cell phylogenies show the amount of changes in methylation in the development of one cell type from another. We show that accounting for CpG island wide methylation changes has a strong effect on the inferred branch lengths and leads to a significantly better model fit for the methylation data from murine haematopoietic cells.

**Availability:** An implementation of the methods presented in this article is freely available as C++ source code on https://github.com//statgenlmu//IWEPoissonPaper under the terms of the GNU general public license (GPLv3).

## 1. Introduction

DNA methylation is a common epigenetic process (Smith and Meissner, 2013). It is considered essential during phenotypic development in mammals and strongly associated with differential gene expression. The most frequent form of methylation is the attachment of the methyl group at the fifth carbon position on cytosine nucleotides that are followed by guanine nucleotide (Saxonov et al., 2006; Eckhardt et al., 2006; Smith and Meissner, 2013). Together, this configuration is called a CpG site.

Regions in which more than 50% of sites are either G or C are called CpG islands if the number of CpGs is greater than 60% of the expected number of CpG sites by random order (Saxonov et al., 2006; Smith and Meissner, 2013). These regions are typically between a few hundred and two thousand base pairs in length (Smith and Meissner, 2013). CpG islands are involved in the regulation of transcription (Deaton and Bird, 2011).

Comparisons of methylation states have been commonly applied and proved as a fruitful avenue of analysis of cell haematopoiesis (Bock et al., 2012; Xie et al., 2013). Pairwise comparison between cell types in different stages of differentiation or comparison between malignant and healthy cells during cancer development have provided insight into areas of transcription (Bock et al., 2012) and enabled inference of missing methylation states. Capra and Kostka (2014) have adapted phylogenetic methods to account for the tree-shaped genealogy of cell types when analyzing methylation changes during haematopoiesis. The branch lengths of the genealogy, representing expected numbers of methylation changes per site, were inferred via likelihood maximization.

A common simplifying assumption in phylogenetics is that sequence positions evolve independently of each other. In an analogous manner, Capra and Kostka (2014) assume that the methylation processes at all CpG sites are, conditioned on the genealogy, stochastically independent of each other. This model assumption is violated when, for example, methylation frequencies change in an entire CpG island in the course of gene regulation (Deaton and Bird, 2011; Smith and Meissner, 2013).

Some sequence evolution models allow that mutation rates change at random time points, see e.g. Huelsenbeck et al. (2000). Here, we adapt this approach to CpG methylation-demethylation dynamics and allow that methylation frequencies in CpG islands can change during island-wide events (called IWEs throughout this text) at random time points. Furthermore, we allow that CpG sites that belong to the same island can simultaneously be methylated or demethylated in an IWE.

We implemented a reversible-jump MCMC inference scheme (Green, 1995; Sorensen and Gianola, 2002; Hastie and Green, 2012) to fit this model to RRBS methylation data (Meissner et al., 2005; Bock et al., 2012). We validate the accuracy of this scheme in a simulation study. With RRBS data procured from mouse haematopoiesis (Bock et al., 2012; Capra and Kostka, 2014) we demonstrate that accounting for IWEs can lead to significantly different estimations of branch lengths of cell type genealogies.

## 2. Methods

### 2.1 Methylation model

We assume that several CpGs can form a CpG islands, which can be affected by CpG island wide events (IWEs) in which methylation probabilities change and some of the CpG sites in the CpG island can simulaneously change their state at this time point. Different CpG islands, however, evolve independently of each other. Following Capra and Kostka (2014) we distinguish three possible states {*u, p, m*} of a CpG site, denoting unmethylated, partially methylated and methylated sites. When analyzing methylation sequencing data (see section 2.5) we consider a site unmethylated if it is methylated in less than 10% of the reads overlapping the site, partially methylated if it is detected as methylated in 10 to 80% of the reads, and methylated if it is methylated in more than 80% of the reads.

For a branch with *h* IWEs set *t*_0_ = 0, let *t*_*h*+1_ be the branch length, and let *t*_1_, …, *t*_*h*_ be the branch-length distances of the IWEs to the parent node. For each CpG position and each open interval (*t*_*k*_, *t*_*k*+1_) there is a rate matrix *Q*_*k*_ for the transitions between the states *u, p, m*. In the open interval (*t*_*k*_, *t*_*k*+1_), the methylation dynamics of CpGs of the same island are independent of each other and the matrix *P*_*k*_ of transition probabilities 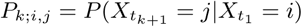 between the methylation states 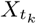 and 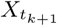 of a CpG at time points *t*_*k*_ and *t*_*k*+1_ can be calculated with the matrix exponential *P*_*k*_ = exp(*Q*_*k*_ *·* (*t*_*k*+1_ *- t*_*k*_)). In analogy to the F81 sequence evolution model (Felsenstein, 1981) we focus here on rate matrices *Q*_*k*_ that can be expressed as

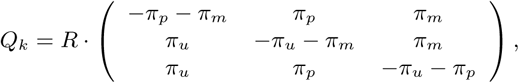

where *π*_*u*_ + *π*_*p*_ + *π*_*m*_ = 1, and each CpG has its own random rate factor *R* ∈ R_*≥*0_. This implies that (*π*_*u*_, *π*_*p*_, *π*_*m*_) is a reversible equilibrium of *Q*_*k*_ and for fixed *R* the transition probabilities (*P*_*k*_)_*i,j*_ take the form 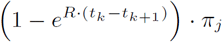 for *j ≠i* and 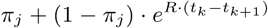. for *j*=*i*. We assume that in the root of the genealogy each CpG island samples a distribution (*π*_*u*_, *π*_*p*_, *π*_*m*_) from a uniform distribution (that is Dirichlet(1,1,1)) independently of all other CpG islands. Like in F81 and related models, the time scaling in our models can be interpreted as follows. At each CpG site with the respective rate *R* events occur that let the CpG sample a new state *u, p* or *m* according to the probabilities (*π*_*u*_, *π*_*p*_, *π*_*m*_). We will refer to these events as single-site events (SSEs) in the following. Note that an expected fraction of 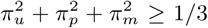 of the SSEs will not change the current state of the CpG. We assume 𝔼*R* = 1, which implies that our time unit is the expected number of SSEs per CpG (not conditioned on *R* but averaged over the possible values of *R*). In the following, branch lengths *B* := (*l*_1_, *l*_2_, …, *l*_*k*_) will refer to this time scaling.

We assume that IWEs occur independently at each CpG island at rate *µ* and change the parameters values *π*_*u*_, *π*_*p*_ and *π*_*m*_ on the CpG island. For a branch of length *l* we obtain an expected number of *l* SSEs per site and of *µ · l* IWEs per CpG island. This implies that if *n* is the number CpG islands and *n*_*i*_ the number of CpG sites on CpG island *i* with 1 *≤ i ≤ n* and the random variables *S* and *W* are the numbers of SSEs and IWEs on a branch of length *l*, we obtain

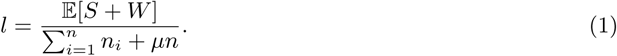

Note here that *S* also counts all SSEs, including those that do not change the methylation state of the site.

In an IWE a new triple of equilibrium methylation frequencies 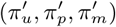 is sampled from a uniform distribution, and *Q*_*t*_ is updated accordingly for time points *t* after the IWE. Furthermore, we allow that CpG sites of an island are methylated or demethylated simultaneously in an IWE in a way such that the expected frequencies of the states *u, p* and *m* match the new equilibrium distribution 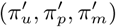 right after the IWE. To specify the transition probability matrix *M*_*k*_ in an IWE at a time point *t*_*k*_ we distinguish two cases. In the first case one of the new expected frequencies is larger and the other two are smaller after the IWE. Without loss of generality, assume 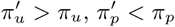 and 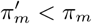 Then the transition matrix is

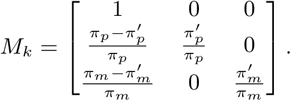

In the other case, one of the new expected frequencies is smaller and both others are larger. If, again w.l.o.g., 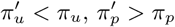 and 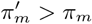, the matrix of transition probabilities is

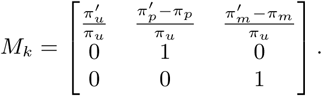

Note that 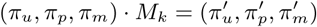 holds in both cases. For given IWEs at time points *t*_1_, …, *t*_*h*_ between time points *t*_0_ and *t*_*h*+1_, the transition matrix between the states {*u, p, m*} at time *t*_0_ and the states at time *t*_*h*+1_ is *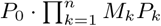*.

For *R* we assume an “invariant+gamma” model (Yang, 1994; Felsenstein, 2004). That is, *R* is 0 with probability *r*, and with probability 1 *− r* the value of *R* comes from a discretized gamma distribution with 3 categories, expectation value 1 and a shape parameter *α*. The probability to be in each respective rate category, conditional on not being an invariant site, is 1/3.

### 2.2 Likelihood calculations

We summarize the global model parameters as *θ* := (*r, α, µ*). As we assume that CpG islands evolve independently of each other, we obtain Pr_*θ,B*_(*D*) = Π_*i*_ Pr_*θ,B*_(*D*_*i*_), where *D*_*i*_ is the data from CpG island *i* and *B* is the vector of branch lengths of the tree. For CpG island *i* let *W*_*i*_ be the configuration of IWEs and the mutation model parameters *π*_*u*_, *π*_*p*_, *π*_*m*_ around them. If we condition on the configuration *W*_*i*_, the CpGs within the island become independent and we obtain Pr_*θ,B*_(*D*_*i*_*|W*_*i*_) = Π_*j*_ Pr_*θ,B*_(*D*_*ij*_*|W*_*i*_), where *D*_*ij*_ is the data from the *j*-th CpG in CpG island *i*. Pr_*θ,B*_(*D*_*ij*_*|W*_*i*_) is a weighted average of Pr_*θ,B*_(*D*_*ij*_*|W*_*i*_, *R*_*ij*_ = *x*), where *x* is iterated over the four possible values of the rate factor *R*_*ij*_ for the CpG position. To calculate Pr_*θ,B*_(*D*_*ij*_*|W*_*i*_, *R*_*ij*_ = *x*) we used a recursive scheme derived from Felsenstein’s pruning algorithm (Felsenstein, 1973, 2004). For this, let 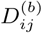 be the part of *D*_*ij*_ that stems from the offspring of branch *b*. For focal *b, i* and *j*, and any *y* ∈ {*u, p, m*} and *k ≥* 1 we now define the partial likelihood *ω*_*k,b*_(*y*) to be the conditional probability of the partial data 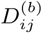, given that CpG site *j* is in methylation state *y* just before IWE *k* (or the child node if *k* = *h* + 1, where *h* is the number of IWEs on *b* affecting island *i*), and given the current states of *θ, B, W*_*i*_ and *R*_*ij*_. For *k ≥* 0 be 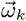 the column vector (*ω*_*k,b*_(*u*), *ω*_*k,b*_(*p*), *ω*_*k,b*_(*m*))^*T*^. Let 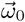 be defined accordingly, but given that the state of CpG *i* in the parent node of the branch *b* is *y*. With the transition probability matrices *P*_*k*_ and *M*_*k*_ as defined in section 2.1 we obtain 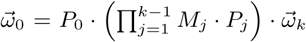 for any *k* ∈ {1, …, *h* + 1}. The case *k* = *h* + 1 is sufficient for likelihood calculations, but the formula is also used for other values of *k* for updating likelihoods when *M*_*k-*1_ and *P*_*k-*1_ are changed in an MCMC step, see online appendix section 2.1.

If the child node of *b* is a tip of the genealogy, we obtain *ω*_*h*+1,*b*_(*y*) = 1 if *y* ∈ {*u, p, m*} is the state of the focal CpG site at the child node, and otherwise *ω*_*m*+1,*b*_(*y*) = 0. If *b* is a node with two daughter nodes *b*^*′*^ and *b*^*″*^, we obtain

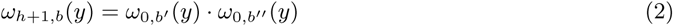

for all *y* ∈ {*u, p, m*}. In our application example below, all methylation states are known not only for the tips of the tree but also for the internal nodes. In this case equation (2) holds only if *y* is the state of the focal CpG site at *b*’s child node, and otherwise *ω*_*h*+1,*b*_(*y*) = 0. For the branch *r* that starts in the root (HSC in our example below) we apply Pr_*θ,B*_(*D*_*ij*_*|W*_*i*_, *R*_*ij*_ = *x*) = *π*_*z,r*_ *· ω*_0,*r*_(*z*), where *z* is the state of the CpG in the root note and *π*_*z,r*_ is its probability according to the equilibrium distribution in the root.

### 2.3 MCMC implementation

To approximate Pr_*θ,B*_(*D*_*i*_) we have to average Pr_*θ,B*_(*D*_*i*_*|W*_*i*_) over possible configurations of *W*_*i*_. For this we apply a Metropolis-Hastings MCMC method (Hastings, 1970; Sorensen and Gianola, 2002). Given the current configuration of *W*_*i*_ in the MCMC procedure, the proposed 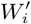 for the next step can either lack one of the IWEs in *W*_*i*_ or have an additional IWE on some branch. Let *l* be the length of a branch *b*. As the IWE locations according to *W*_*i*_ are *a priori* a Poisson point process with intensity *µ*, the prior probability that *W*_*i*_ includes *n* IWEs on branch *b* is Pois_*µ·l*_(*n*) = (*µl*) ^*n*^ *e*^− *µl*^ /*n*!. When the proposed 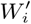 differs from the current *W*_*i*_ by an additional IWE on branch *b* and *n* is the current number of IWEs on this branch, the Metropolis-Hastings acceptance probability is the minimum of 1 and

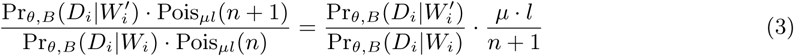

(see online appendix 2.1). If, conversely, 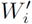 with *n* + 1 IWEs on *b* is the current state, and *W*_*i*_ with one IWE less on *b* is proposed, the acceptance probability is the minimum of 1 and the inverse of any side of equation 3.

For the branch lengths we apply Metropolis-Hastings acceptance steps on the log scale. If *ℓ* = log(*l*) is the (natural) logarithm of the current length of a branch, the proposed *ℓ*^*′*^ = log(*l*^*′*^) is drawn from a Gaussian mixture proposal distribution with density *g*_*.ℓ*_(*ℓ*^*′*^) that is centered around *ℓ*. The proposal distribution is symmetric, that is *g*_*.ℓ*_(*ℓ*^*′*^) = *g*_*.ℓ′*_ (*ℓ*). The prior distribution of *ℓ* is a normal distribution. Let *p*(*ℓ*) denote its density (see online appendix 1). When a new log length *ℓ*^*′*^ is proposed for a branch of log length *ℓ*, we obtain the acceptance probability

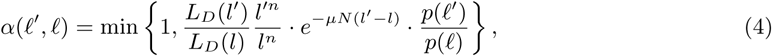

where *n* is the current number of IWEs on the branch, *µ* is the rate of IWEs, *N* is the number of CpG islands, *L*_*D*_ (*l*^*′*^) and *L*_*D*_ (*l*) are the conditional probabilities of the data given the proposed and the current trees.

In further Metropolis-Hastings steps, the methylation state frequencies (*π*_*u*_, *π*_*p*_, *π*_*m*_) for any CpG island can be updated. Further information about priors and proposal densities can be found in the online appendix 1.

### 2.4 Null model without IWEs

We test our model against a null model without IWEs. In the null model we still assume that each island has distinct equilibrium frequencies, which are sampled at the root from a Dirichlet(1,1,1) distribution and do not change during sequence evolution. When new branch lengths are sampled, the acceptance probability (4) in this model simplifies to

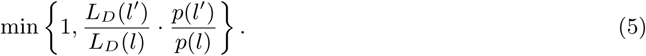

The parameters of the null model are the logarithms of branch lengths, the logarithm of the shape parameter of the gamma distribution of site specific rate factors, the fraction of invariant sites, and for each CpG island the equilibrium probabilities at the root states.

### 2.5 Data

We tested our approach with methylation data that were gained by Bock et al. (2012) with reduced restricted bisulfite sequencing (RRBS) from murine cells at various stages of haematopoiesis (Figure 1). The data overlap most murine CpG islands and consist of reads that are 36 base pairs long. To associate information of reads with CpG islands we used the mmp9 mapping of CpG islands from the UCL genome browser (Kent et al., 2002). We sampled 2000 CpG islands at random, 1970 of which contained reads overlapping CpGs within the island. CpGs were categorized as unmethylated (*u*), partially methylated (*p*) or methylated (*m*) if less then 0.1, between 0.1 and 0.8, or more than 0.8 of the reads were detected as methylated. For Fig. 2 we categorized whole CpG islands as unmethylated if more than 50 % of its CpG sites were in state *u*, or as methylated if if more than 50 % of its CpG sites were *m*. All other CpG islands were classified as partially methylated.

**Figure 1:**
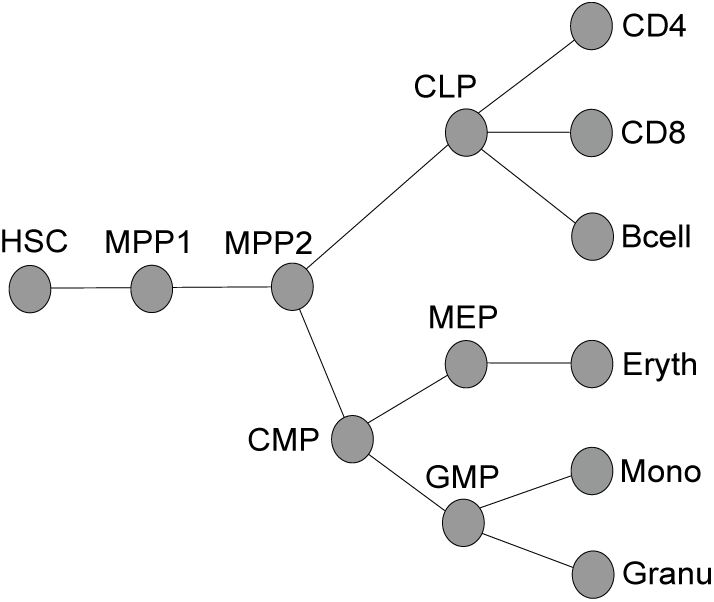
Genealogy of haematopoietic cell stages (Bock et al., 2012; Capra and Kostka, 2014).

**Figure 2:**
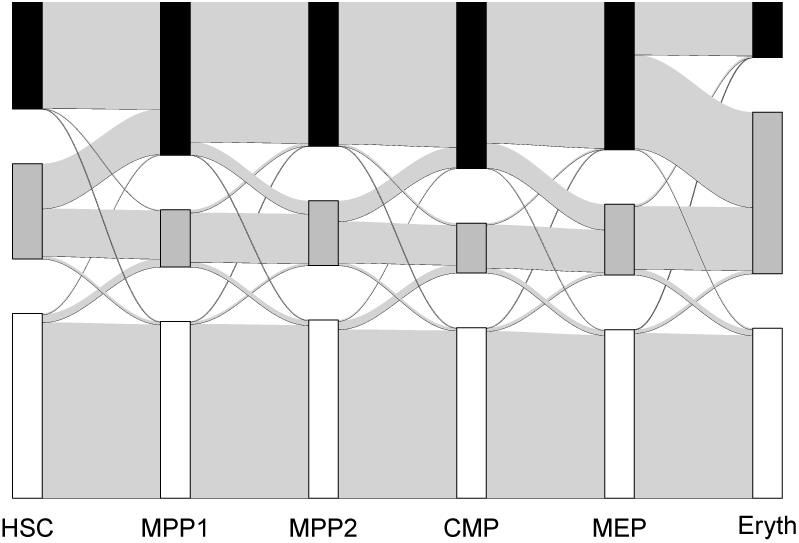
Change of Island Methylation States. States of islands are categorized as methylated (black, top) if more than 50% of sites are in state *m* and as unmenthylated (white, bottom) if more than 50% of sites are in state *u*. Otherwise, islands are categorized as partially methylated (grey, middle). Vertical rectangles are proportional in size to the respective number of islands in each state for each cell type. Light grey transitions have a width proportional to the relative amount islands that transition between the states indicated by the rectangles.

### 2.6 Simulation study

We assess the accuracy of our MCMC implementation in a simulation study. We simulated 150 data sets, each consisting of 100 islands. The number of CpG sites in each islands were chosen randomly from a uniform distribution between 10 and 400. At the start of each simulation we sampled the logarithm of branch lengths, the logarithm of the shape parameter *α*, the invariant probability, and the IWE rate *µ* from our priors. For the root node we sampled equilibrium frequencies from a Dirichlet(1,1,1) distribution. Then we sampled IWEs uniformly positioned along branches, where the number of sampled IWEs on a branch was Poisson distributed with mean *µNl*, where *N* is the number of CpG islands and *l* is the branch length. The equilibrium frequencies associated with an IWE were sampled from a Dirichlet(1,1,1) distribution. Given a vector of equilibrium frequencies and positions for IWEs occurring at an island we calculated transition probabilities between states as detailed in the model specifications.

We generated the sequence at the root node by drawing each state in each island from the equilibrium frequency at the root node in this island. Sequences in the other nodes where generated iteratively going from the root to the tips of the cell lineage tree.

We then used our inference method on the generated sequences to find posterior distributions of the simulated data sets with known ground truths sampled from priors. Here the MCMC runs were started from the means of the priors for all parameters other than the number of IWEs, where we started without IWEs to avoid long convergence times in the case of many misplaced IWEs in the initial configuration. We used a burn-in of 10^5^ Metropolis Hasting steps.

### 2.7 Test for CpG-island-wide events (IWEs)

In addition to our full model we also fitted a null model without IWEs to the data. To test the relevance of IWEs for the data of Bock et al. (2012), we simulated 150 data sets according to this null model using 1970 islands with the same number of CpG sites as in the restricted data set we used for initial inference. These simulations were conducted with the same procedure as in the simulation study, with the starting parameters being sampled from the posterior distribution of the null model and the IWE rate being restricted to 0. We then fitted the full model with IWEs to these simulated sequences and estimated posterior number of IWEs inferred in the adapted model.

## 3. Results

### 3.1 Application to methylation data from haematopoietic cells

We applied our method to 1970 randomly chosen CpG islands from the methylation data of murine haematopoitic cells (Bock et al., 2012; Capra and Kostka, 2014). Using first the null model without IWEs (*µ* = 0), we obtained a very long branch between MEP and Eryth indicating many changes in methylation of single CpGs (Fig. 3). Note that branch lengths are proportional to the expected number of changes of events on the branch or, in other words, to the product of cell divisions and the rate of methylation and demethylation per cell division. We generated data following this fitted null model in 150 simulations and estimated parameters according to the model with IWEs for the null model simulations. Estimated total numbers of IWEs never exceeded 30 in any of these inferences and the inferred percentage of islands carrying an IWE along an edge was a most 0.07%. When we analysed the data set of Bock et al. (2012) with the IWE model (*µ* ≥ 0), the minimum number of IWEs after the burn-in period of 10^6^ Metropolis-Hasting steps was 3488, and we inferred high levels of enrichment of IWEs on all branches (Fig. 4).

**Figure 3:**
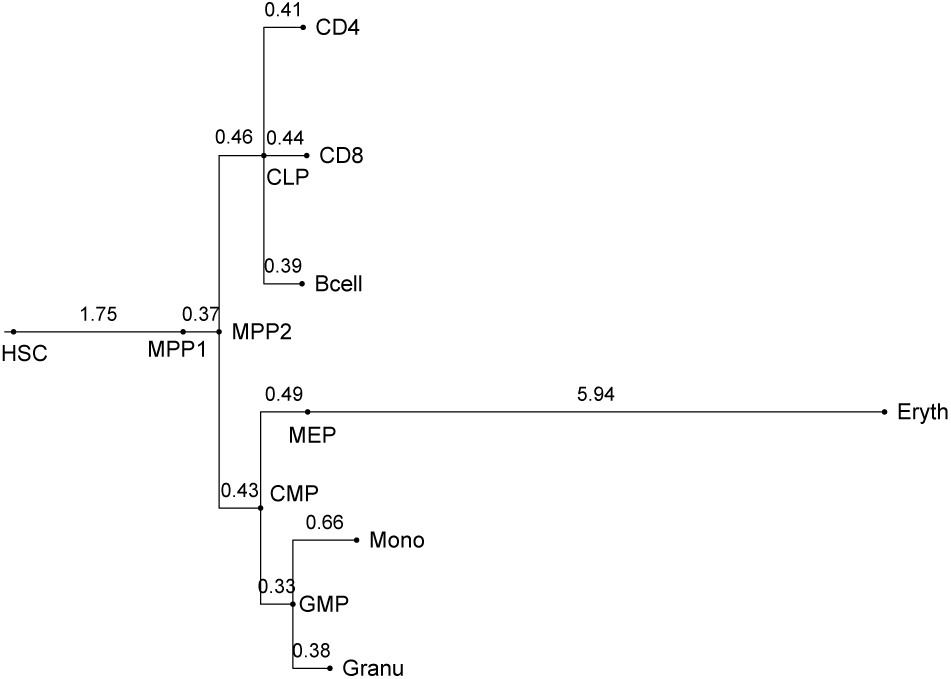
Tree resulting from estimates without modeling IWEs. The logarithmic branch lengths are the means of the MCMC samples after a burn in phase of 10^6^ steps.

**Figure 4:**
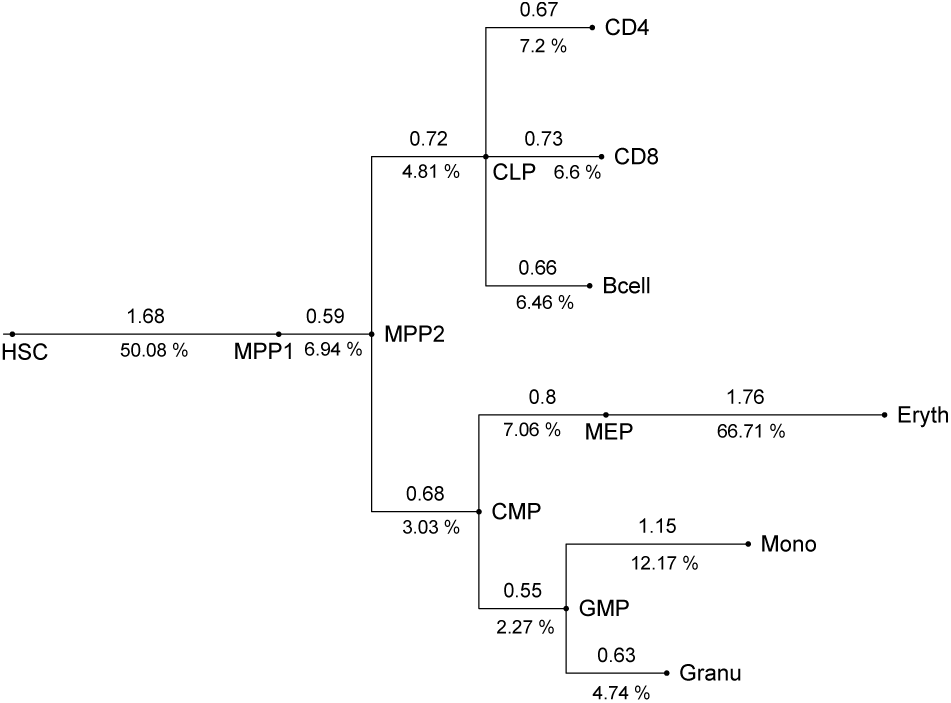
Tree resulting from estimates modeling IWEs. The logarithmic branch lengths (above branches) and percentages of CpG islands affected by IWEs (below branches) are the means of the MCMC samples after a burn in phase of 10^6^ steps.

With the null model we estimated branch lengths similar to estimated lengths in the literature on these branches, e.g. between MEP and Eryth 4.56 units by Capra and Kostka, compared to a distribution mean of 5.94 SSE units with our null model. Here, an SSE unit refers to the expected number of SSEs per CpG, whereas Capra an Kostka’s unit refers to the expected number of methylation state changes per CpG. As at least a third of the SSEs do not change the state of a CpG, and 5.94 *·* 2*′*3 = 3.96, our estimation of the length of the MEP-Eryth branch is smaller than that of Capra and Kostka, but the values are not directly comparable, because the model of Capra and Kostka is more general than our null model without IWEs.

When we allowed for IWE events, we found considerably less variation among the inferred branch lengths (Fig. 4). Regarding the number of IWEs, the formation of the first multi-pluripotent cells from haematopoietic stem cells and the formation of erythrocytes showed an increased frequency of such events, explaining the methylation changes between MEP and Eryth by simultaneous methylation changes in IWEs rather than by many independent single-site events.

#### 3.1.1 Evidence that IWE rate vary among branches

In the tree that we inferred with the IWE model (Fig. 4), the estimated numbers of IWEs vary among the branches more than the branch lengths. Indeed, credibility intervals of the log-transformed numbers of IWEs per branch length unit (Fig. 5) suggest that the IWE rate is substantially increased during the transitions from HSC to MPP1, from MPP1 to MPP2 and from MEP to Eryth. This is indicative of pronounced regularly activity along these transitions in particular (see also Fig. 2).

**Figure 5:**
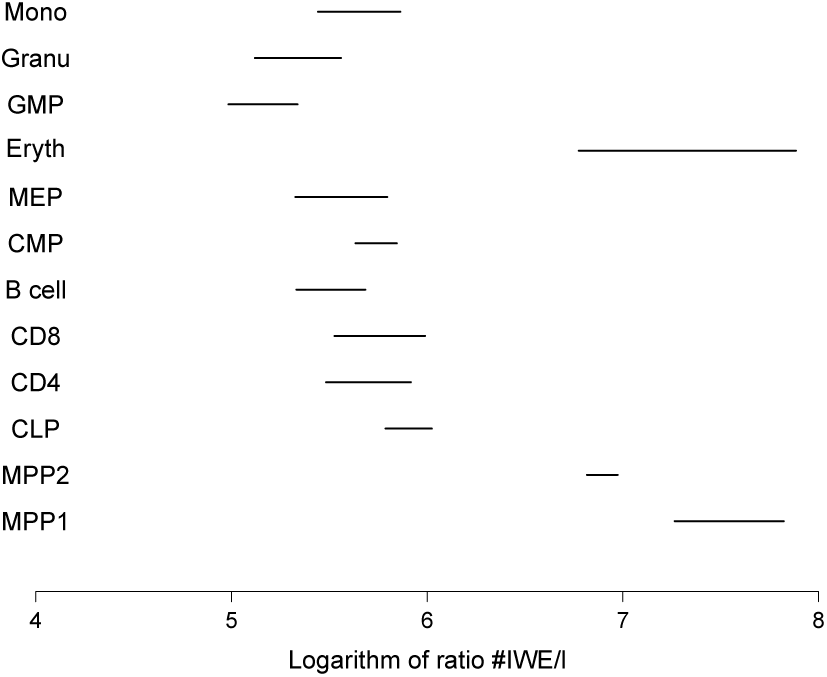
Multiple testing corrected 95% intervals of the ratio of estimated number of events to the estimated branch length.

### 3.2 Simulation experiments

To validate the accuracy of our inference we conducted, we simulated 150 data sets with parameters values drawn from the prior distributions (see 2.3 and online appendix 1). Each of the simulated data sets contained 100 islands with sizes varying uniformly between 10 and 400. For each of the simulated data sets we inferred the posterior distribution of the parameters we used to produce the simulated data. In Fig. 6 we compare the MCMC-sampled parameter values and credibility intervals to the actual parameter values underlying the simulations. To validate our implementation we computed the 95% credibility intervals and verified that the ground truth was within the these intervals in approximately 95% of the cases. This was done for individual branch lengths, all branch lengths, the rate of IWEs and the shape parameter of rate heterogeneity. Indeed, credibility intervals overlapped the true branch length in 93 to 98% of the cases. Overall, 95% of the credibility intervals contained the true value. True values were in the credibility intervals in 96 % of the cases for IWE rates and in 94 % for the shape parameter.

**Figure 6:**
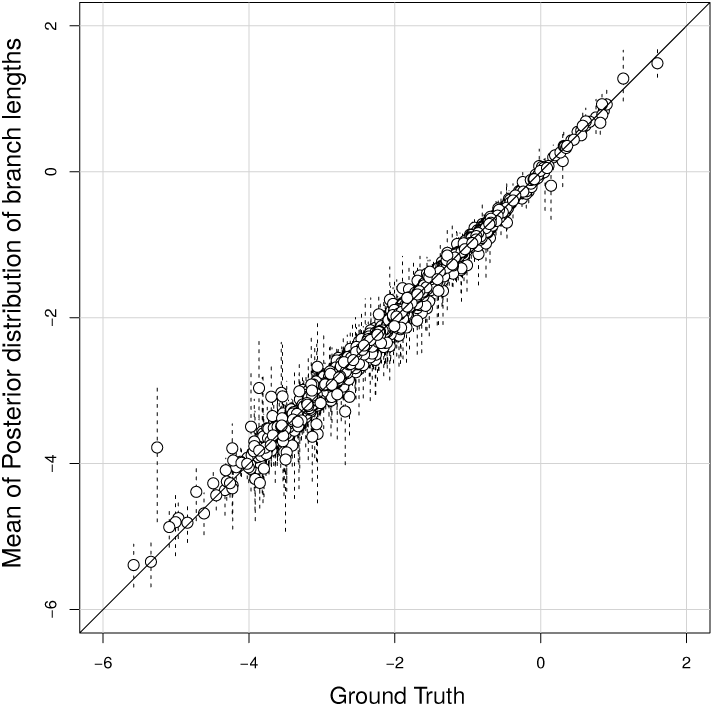
Comparison between estimated logarithms of branch lengths and target values in simulation study. Dotted lines indicate 95% credibility intervals.

## 4. Discussion

We found that the model with CpG-island wide methylation rate changes (IWEs) fit the methylation data from murine haematopoietic cells significantly better than a model without IWEs. Furthermore, the IWE model suggested for certain developmental phases in haematopoiesis that many CpG islands were affected by IWEs, which may indicate enhanced activity in gene regulation.

For the single-site methylation changes (SSEs) we assume in our current model that the new methylation state (unmethylated, partially methylated, or methylated) is independent of the state before the SSE. A possible extension of our model would be to allow for the SSEs and for state-changes within IWEs the class of models proposed by Capra and Kostka (2014), who consider all reversible 3 *×* 3 rate matrices for the three states.

Even though we assumed *a priori* a constant IWE rate in our model, we obtained clear evidence that the number of IWEs per branch length unit (which summarizes expected numbers of IWEs and SSEs) varies among the branches (Fig. 5). Further, Figure 2 suggests that overall methylation frequencies vary among the branches. Also this is not explicitly taken into account in our model, as we assume that IWEs have their probabilities sampled from the same Dirichlet distribution across the tree. However, compound Poisson based models (Huelsenbeck et al., 2000) of genome wide change are natural extensions to our framework. Thus, we could allow for genome-wide events that modify the IWE rate and the parameters of the Dirichlet distribution from which the methylation state distribution are sampled in IWEs. An alternative approach, in analogy to some relaxed molecular-clock models in phylogenetics (Drummond et al., 2006), would be to assume that IWE rates or other parameters are sampled from a prior distribution independently for each branch.

In the application example above with the data of Bock et al. (2012), the tree topology and the methylation states at the internal nodes were given. Our computational approach for the IWE-SSE model can also be adapted to reconstruct genealogies when methylation states are given only for the tips of the tree and combined with methods to explore possible tree topologies (Felsenstein, 2004). A potential application area could then be the inference of genealogies of cells sampled from neoplasms, e.g. to reconstruct the growth and mutation history of cancer clones (Brocks et al., 2014; Sottoriva et al., 2013). Accounting for IWEs may not only improve the accuracy of inferred cell genealogies but also allow for a better detection of aberrant methylations, which are known to be among the hallmarks of cancer (Hanahan and Weinberg, 2011). The best possible data for reconstructing cell genealogies from methylation patterns would obviously be single-cell methylation data. To our knowledge, however, it is not yet possible to generate such data. Therefore it seems worthwhile to explore possibilities of inferring single cell genealogies from long-read methylation data, which are now becoming available (Simpson et al., 2017).

## Acknowledgments

We thank Tobias Altmiks for software testing an for spotting a bug in an early version of our program code.

