## Supplemental Material for "Modeling methylation dynamics with simultaneous changes in CpG islands"

### ONLINE APPENDIX FOR
### Modeling methylation dynamics with simultaneous changes in CpG islands

Konrad Grosser and Dirk Metzler
Faculty of Biology, Division of Evolutionary Biology
Ludwig-Maximilians-Universität München

May 14, 2019

#### 1 Priors and proposal densities for Parameters

In several types of Metropolis-Hastings steps we use Gaussian mixture proposal densities, which simplify Metropolis-Hasting acceptance ratios to Metropolis acceptance ratios due to symmetry (Metropolis et al., 1953; Hastings, 1970). These proposals are mixtures of two Gaussian distributions $\tilde{g}_x$ and $\tilde{g}'_x$ with mean being the current value $x$ and different standard deviations. With a probability of 0.5 the new proposal $x'$ has probability density $\tilde{g}_x(x')$, and else $\tilde{g}'_x(x')$.

##### 1.1 Estimating a new IWE rate

We estimate the logarithm of the IWE rate $\mu$. As prior we used a normal distribution with mean $-2$ and standard deviation of 1. The Gaussian mixture proposal probabilities for $\log(\mu)$ were normally distributed with a standard deviation of 1 or 0.2, each of them chosen with 50% probability. When computing the acceptance probability $a\left(\log(\mu)', \log(\mu)\right)$, not the whole conditional probability of the data needed to be computed. In particular the probability of the data given the tree and IWEs on the trees does not change, and hence

$$a\left(\log(\mu)', \log(\mu)\right) = \min\left\{1, \frac{P\left(D|\mu', T\right)}{P\left(D|\mu, T\right)} \cdot \frac{\pi(\log(\mu'))}{\pi(\log(\mu))}\right\} = \min\left\{1, \frac{P\left(D|T\right) \cdot P(T|\mu')}{P\left(D|T\right) \cdot P(T|\mu)} \cdot \frac{\pi(\log(\mu'))}{\pi(\log(\mu))}\right\}, \quad (1)$$

where $T$ denotes branch lengths, number and location of IWEs and associated probabilities and probabilities at the root. Terms deriving from the proposal probability are not considered because the proposal is symmetric. Since $T$ only depends on $\mu$ via the number of IWEs in the tree, and the number of IWEs in the tree is Poisson distributed with rate $\mu \cdot L \cdot N$, $L$ being the length of the tree and $N$ being the number of CpG islands, we obtain

$$a\left(\log(\mu)', \log(\mu)\right) = \min\left\{1, \frac{\mu'^n}{\mu^n} \cdot \exp(-L \cdot N \cdot (\mu' - \mu)) \cdot \frac{\pi(\log(\mu'))}{\pi(\log(\mu))}\right\}, \quad (2)$$

where $n$ is the number of IWEs.

##### 1.2 Estimating a new shape parameter

For the logarithm of the shape parameter $\alpha$ we used a normal prior of mean 2 and standard deviation 1. Proposal probabilities for the logarithm of $\alpha$ were normally distributed with a standard deviation of 0.1. When determining acceptance of a newly proposed shape parameter, the conditional probability of the data had to be recalculated. Terms from the proposal probability do not have to be considered due to symmetry.

##### 1.3 Estimating a new invariant probability

For the invariant probabilities we used a uniform prior, and proposal probabilities were normally distributed and are reflected on the boundaries of the interval $[0, 1]$. The standard deviation of the normal distribution was 0.1. When estimating a new invariant probability the whole conditional probability of the data given $T$ had to be updated.

##### 1.4 Other priors and relative frequency of steps

Logarithms of branch lengths had a normal prior with mean $-2$ and standard deviation 1. Proposals for adding an IWE and deleting an IWE (if IWE existed) happened with probability 0.5 each.

The relative frequency of proposing new branch lengths, sampling new shape parameters, sampling new $\mu$, sampling or deleting an IWE, sampling an invariant probability, and sampling a new beginning frequency in an island were $0.1, 0.01, 0.1, 1, 0.01$ and 0.3, respectively.

#### 2 Deriving acceptance probabilities

When new IWEs are inserted on a branch, the parameter space from which our MCMC procedure samples is augmented by the location of the IWE on the branch and by the new methylation state probabilities. The varying dimensionality of the MCMC state space is handled by reversible jump MCMC (Green, 1995). Here, we follow the formalism of Hastie and Green (2012), in which the proposal to move from state $x$ to state $x'$ is constructed with an additional random vector $u$ with probability density $g(u)$ and a (deterministic) diffeomorphism $h : \mathcal{U} \times \mathcal{V} \to \mathbb{R}^{n'} \times \mathbb{R}^{r'}$, $(x, u) \mapsto (x', u')$ with $\mathcal{U} \subseteq \mathbb{R}^n$, $\mathcal{V} \subseteq \mathbb{R}^r$ and $n + r = n' + r'$. The Metropolis-Hastings acceptance probability is then

$$a(x, x') = \min \left\{ 1, \frac{P(x') \cdot g'(u')}{P(x) \cdot g(u)} \cdot \left| \det \left( \frac{\partial h(x, u)}{\partial (x, u)} \right) \right| \right\}, \tag{3}$$

where $P(.)$ is the target probability density (usually the posterior), and $g'(u')$ is the proposal density of $u'$ if the current state is $x'$ (Hastie and Green, 2012).

##### 2.1 Deriving transition probabilities for number of IWEs

We are sampling new IWEs and deleting existing IWEs according to the following procedure. For this, we first decide with a fair coin flip whether a deletion or an insertion shall be proposed.

- If insertion, do the following:
  1. Pick a branch with probability proportional to the branch length.
  2. Pick an island at random.
  3. An event is inserted at a random location on that island while the rest remains the same.
  4. Calculate acceptance probability and determine whether we accept.
  5. Revert the tree if not accepted, or else proceed.
- If deletion, do the following:
  1. Pick an IWE at random and delete it. (If no events present, do nothing).
  2. Calculate acceptance probability and determine whether we accept.
  3. Revert the tree if not accepted, or else proceed.

The acceptance probability for an insertion is then

$$\min \left\{ 1, \frac{\mu N t}{n+1} \frac{L_{n+1}}{L_n} \right\}, \tag{4}$$

where $t$ is the total length of the tree $n$ is the number of IWEs in the tree, $\mu$ is the IWE rate, $N$ is the number of islands and $L_n$ and $L_{n+1}$ are the conditional probabilities of the data given the state with $n$ events and the state with $n+1$ events respectively.

The acceptance probability of a deletion on the other hand is

$$\min\left\{1, \frac{n+1}{\mu N t}\frac{L_n}{L_{n+1}}\right\}. \tag{5}$$

This probability is derived following the proof of Huelsenbeck et al. (2000) of a similar proposition for a compound Poisson process used for relaxing molecular clock assumptions in phylogenetics. First we define $T = N \cdot t$ which is the total length of the tree counting all islands separately. Our model requires that each event has a uniform position along the tree, which is equivalent to having a uniform position along $[0, T]$, when we regard the tree as constant and only treat steps concerning the deletion and insertion of IWEs. We can a state $x$ as a vector of the form

$$x = (n, t_1, \pi_{u1}, \pi_{p1}, \pi_{m1}, \ldots, t_n, \pi_{un}, \pi_{pn}, \pi_{mn}), \tag{6}$$

where $t_i$ is the time of occurrence for event $i$ and $(\pi_{ui}, \pi_{pi}, \pi_{mi})$ are the equilibrium frequencies of that event and $n$ is the number of events present. The density of $t_i$ is a uniform density on $[0, T]$, the triples $(\pi_{ui}, \pi_{pi}, \pi_{mi})$ follow a Dirichlet(1,1,1) distribution. $n$ in our model is Poisson distributed with mean $T\mu$. Hence the conditional prior density is

$$\pi(x) = \frac{(T\mu)^n}{n!}e^{-T\mu}\frac{1}{T^n}\frac{2^n}{\sqrt{3}^n}. \tag{7}$$

We write the other state $x'$ as

$$\begin{aligned} x' \quad = \quad & (n+1, t_1, \pi_{u1}, \pi_{p1}, \pi_{m1}, \ldots, t_{k-1}, \pi_{u(k-1)}, \pi_{p(k-1)}, \pi_{m(k-1)}, \\ & t_{\text{new}}, \pi_{u\,\text{new}}, \pi_{p\,\text{new}}, \pi_{m\,\text{new}}, \\ & t_k, \pi_{uk}, \pi_{pk}, \pi_{mj}, \ldots, t_n, \pi_{un}, \pi_{pn}, \pi_{mn}). \end{aligned}$$

Note here that the position where the new elements were inserted were between element $k-1$ and $k$. We call this move $m$ and the probability of inserting at this position in the vector is uniformly $j_m(x') = 1/2(n+1)$. The reverse probability of deleting one of the $n+1$ events is again $j_m(x) = 1/2(n+1)$, which corresponds to taking one of the IWEs at random and deleting it. Note here that the two move types are exactly inverse to each other: Either we insert at position $j$ into the vector or we delete at this position, instead of for example always inserting at the back, which would make deletion not occurring at the back not a reversible move. The factor of 2 comes from deciding whether we delete or accept an event with a fair coin. The conditional prior density is, analogously to $x$

$$\pi(x') = \frac{(T\mu)^{n+1}}{(n+1)!} \cdot e^{-T\mu} \cdot \frac{1}{T^{n+1}} \cdot \frac{2^{n+1}}{\sqrt{3}^{n+1}}. \tag{8}$$

Following Hastie and Green (2012), we construct the dimension jump from $x$ to $x'$ with a random vector $u = (u_t, u_u, u_p, u_m)$, in which $u_t$ follows an uniform distribution on $[0, T]$, while the triple $u_u, u_p, u_m$ follows a Dirichlet(1,1,1) distribution. This leads to the joint density of

$$g(u) = \frac{2}{\sqrt{3}T}. \tag{9}$$

This density defines the probability of a given set on $[0, T] \times \Delta_3$ as the product of the uniform densitis on $[0, T]$ and on the 3-Simplex $\Delta_3$, which is equivalent to a proposal distribution where the time of a new event is sampled uniformly along the tree and the probabilities corresponding to the event are Dirichlet(1,1,1) distributed and sampled independently from the position of the new event. This contribution of the proposal density is in addition to the contribution of the move type discussed above. The corresponding random vector $u'$ when moving from $n+1$ events to $n$ events is just a length zero vector with probability of 1, due to the fact that no new parameters are sampled and the proposition only depends on the move type determining which event is deleted. Define

$$\begin{array}{rcl} h: \quad \{n\} \times \prod_i^{n+1}([0,T]\times\Delta_3) & \to & \{n+1\}\times\prod_i^{n+1}([0,T]\times\Delta_3) \\ (n, t_1, \pi_{u1}, \pi_{p1}, \pi_{m1}, \ldots, t_n, \pi_{un}, \pi_{pn}, \pi_{mn}, u_t, u_u, u_p, u_m) & \mapsto & (n+1, t_1, \pi_{u1}, \pi_{p1}, \pi_{m1}, \ldots, \\ & & t_{j-1}, \pi_{u(j-1)}, \pi_{p(j-1)}, \pi_{m(j-1)}, \\ & & u_t, u_u, u_p, u_m, \\ & & t_j, \pi_{uj}, \pi_{pj}, \pi_{mj}, \ldots, t_n, \pi_{un}, \pi_{pn}, \pi_{mn}). \end{array}$$

Note that $h$ is a permutation. Hence it is a diffeomorphism and the absolute value of the determinant of its Jacobian is 1. Thus, the acceptance probability for the proposed step is

$$\min\left\{1, \frac{\pi(x')}{\pi(x)} \cdot \frac{L(x')}{L(x)} \cdot \frac{j_m(x')}{j_m(x)} \cdot \frac{g'(u')}{g(u)}\right\}. \tag{10}$$

Inserting the distributions we have described exactly produces the acceptance probability of insertion discussed above. The acceptance probability of deletion can be derived analogously.

#### 2.2 Acceptance probabilities for branch length changes

When a branch length is changed, the dimensionality of the state does not change, but its size does. In this case we can also apply the formalism of Hastie and Green (2012) to derive the acceptance probability. We operate on log scaled branch lengths. The proposal is normally distributed with mean at $\log(l)$ and standard deviation $\sigma$. In our example $x = (x_1, \ldots, x_n, \log(l))$, $x' = (x'_1, \ldots, x'_n, \log(l'))$, and the random variable $u$ is one-dimensional as e we are only sampling $\log(l)$ and adjust the other parameters according to the newly sampled length. We assume $g$ to be normally distributed with mean 0 and standard deviation $\sigma$, and $g' = g$. We define

$$\begin{array}{c} h: \mathbb{R}_+^{n+1}\times\mathbb{R} \to \mathbb{R}_+^{n+1}\times\mathbb{R} \\ (x_1, \ldots, x_n, \log(l), u) \mapsto \left(\frac{\exp(\log(l)+u)}{l}x_1, \ldots, \frac{\exp(\log(l)+u)}{l}x_n, \log(l)+u, u\right) \end{array} \tag{11}$$

This produces our wished proposal density for $\log(l')$. The probability that $\log(l') > L$ is the probability that $u + \log(l) > L$ which means that the proposal density derived from $g$ is normally distributed with mean $\log(l)$, which is the proposal density we are using. To calculate $a$ note that $u' = u$ and $g' = g$, so the proposal does not exert influence. We now need to evaluate the determinant of the Jacobian. Clearly

$$\begin{array}{rcl} \partial x'_i/\partial x_j & = & \frac{l'}{l}\delta_{i,j} \\ \partial \log(l)'/\partial \log(l) & = & 1 \\ \partial u'/\partial \log(l) & = & 0 \\ \partial u'/\partial u & = & 1. \end{array}$$

Hence the Jacobian is an upper triangular matrix, whose determinant is the product of the diagonal entries. This yields

$$\det\left(\frac{\partial h(x,u)}{\partial(x,u)}\right) = \frac{l'^n}{l^n}. \tag{12}$$

### 3 Simulation Study Figures

The following figures show the mean of the posterior distribution and the 95%credibility intervals plotted against the ground truth.

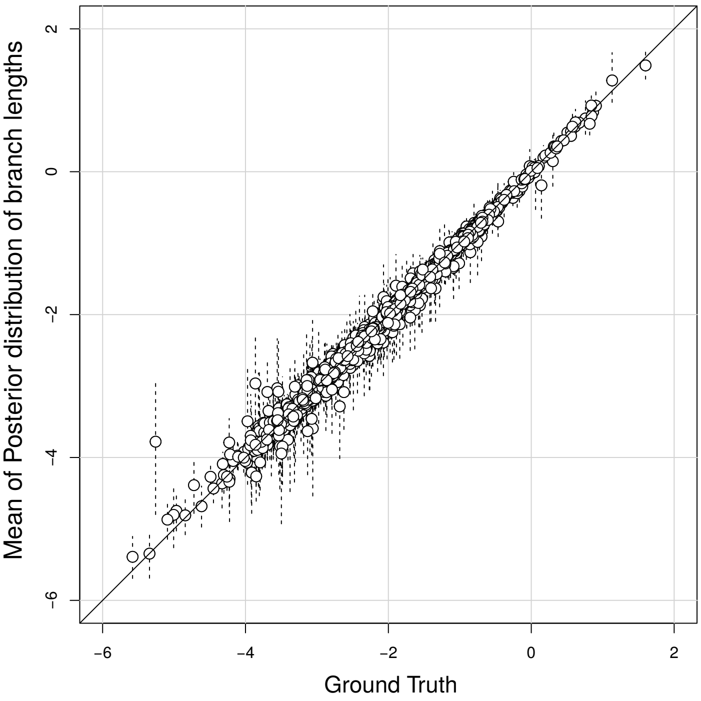


Figure 1: This plot shows all 1800 estimated logarithmic branch lengths compared to the respective actual values used to produce the sequences we infer upon. Black line is the identity. Dotted lines indicate the 95% credibility interval. The true value is expected to fall within in this interval 95% of the time.

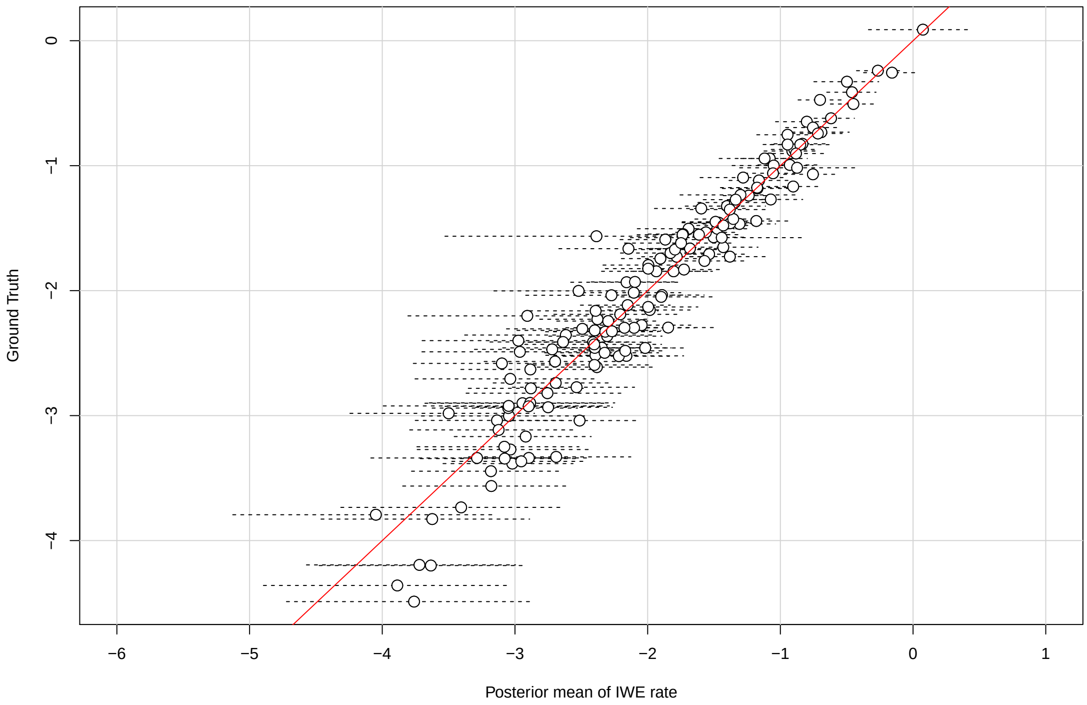


Figure 2: All 150 estimated logarithmic IWE rates compared to the respective actual values used to produce the sequences we infer upon. Red line is the identity. Dotted lines indicate the 95% credibility interval. The true value is expected to fall within in this interval 95% of the time.

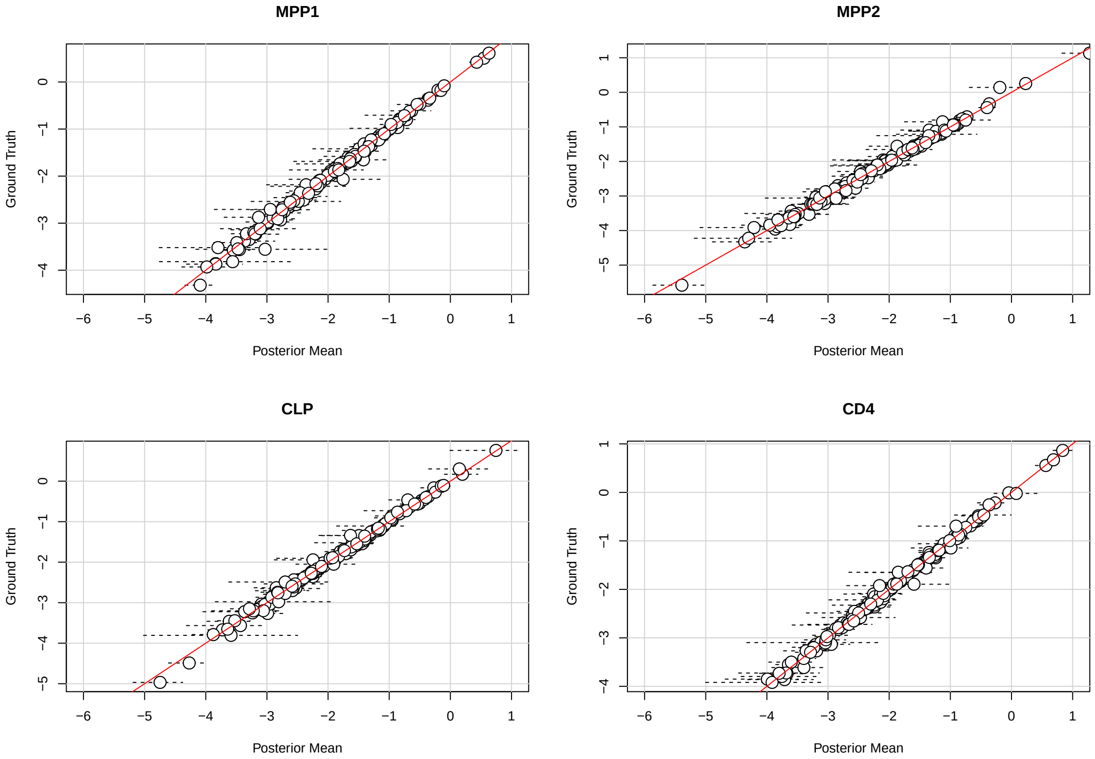


Figure 3: 150 estimated logarithmic branch lengths for the formation of MPP1, MPP2, CLP and CD4 respectively, compared to the actual values used to produce the sequences we infer upon. The true value is expected to fall within in this interval 95% of the time. Red line is the identity.

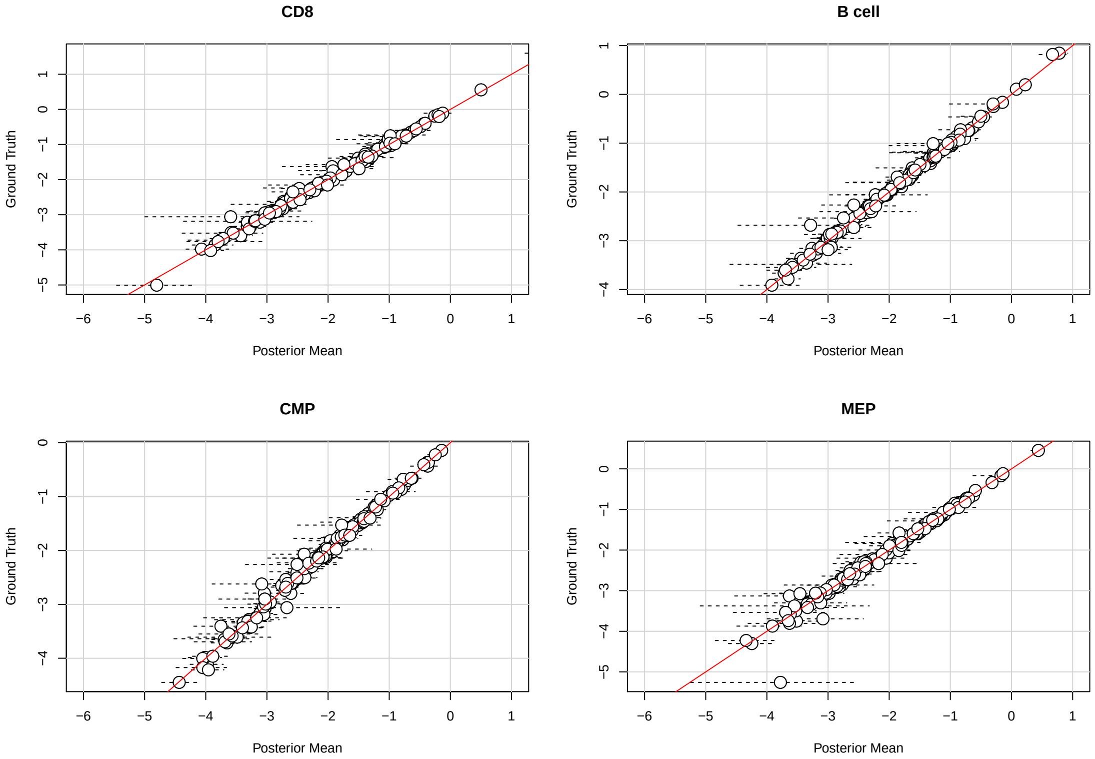


Figure 4: 150 estimated logarithmic branch lengths for the formation of CD8, Bcell, CMP and MEP respectively, compared to the actual values used to produce the sequences we infer upon. The true value is expected to fall within in this interval 95% of the time. Red line is the identity.

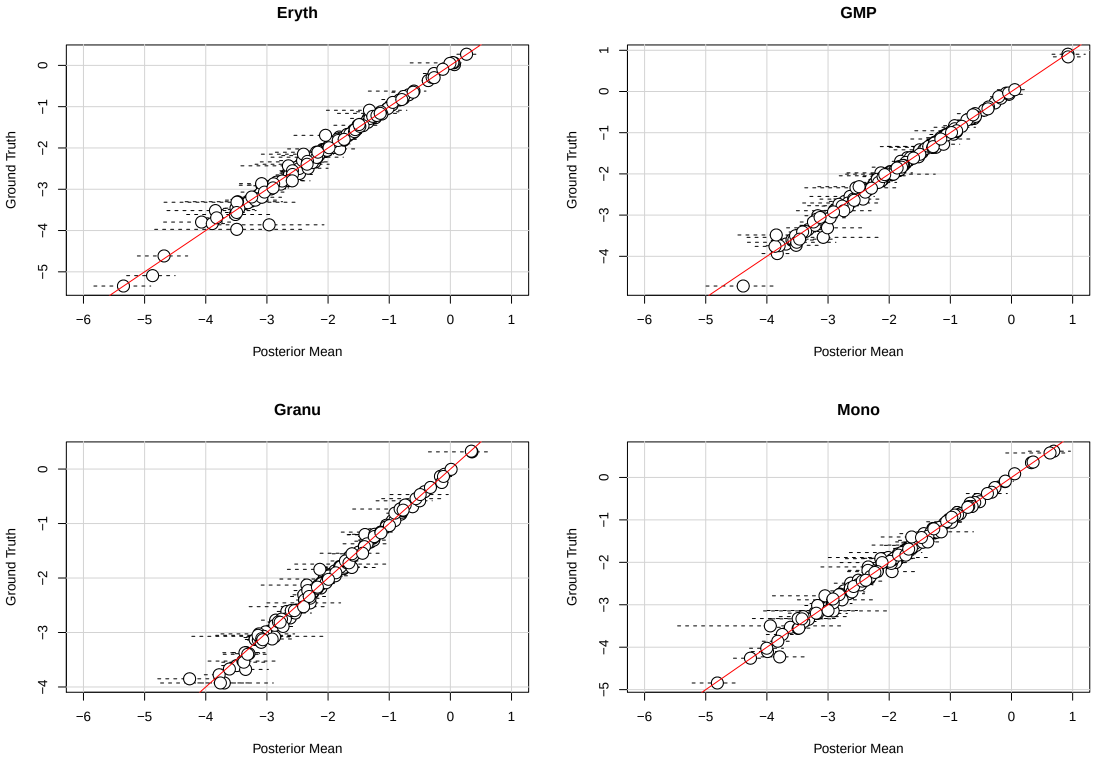


Figure 5: 150 estimated logarithmic branch lengths for the formation of Eryth, GMP, Granu and Mono respectively, compared to the actual values used to produce the sequences we infer upon. Dotted lines indicate the 95% credibility interval. The true value is expected to fall within in this interval 95% of the time. Red line is the identity.
